# Social prediction modulates activity of macaque superior temporal cortex

**DOI:** 10.1101/2021.01.22.427803

**Authors:** Lea Roumazeilles, Matthias Schurz, Mathilde Lojkiewiez, Lennart Verhagen, Urs Schüffelgen, Kevin Marche, Ali Mahmoodi, Andrew Emberton, Kelly Simpson, Olivier Joly, Mehdi Khamassi, Matthew FS. Rushworth, Rogier B. Mars, Jérôme Sallet

**Affiliations:** Wellcome Centre for Integrative Neuroimaging, Department of Experimental Psychology, University of Oxford, Oxford, United Kingdom; Institute of Psychology, University of Innsbruck, Innsbruck, Austria; Wellcome Centre for Integrative Neuroimaging Centre for Functional MRI of the Brain (FMRIB), Nuffield Department of Clinical Neurosciences, John Radcliffe Hospital, University of Oxford, Oxford, United Kingdom; Donders Institute for Brain, Cognition and Behaviour, Radboud University Nijmegen, Nijmegen, The Netherlands; Biomedical Sciences Services, University of Oxford, Oxford, United Kingdom; Institute of Intelligent Systems and Robotics, Sorbonne Université, CNRS, Paris, France; Univ Lyon, Université Lyon 1, Inserm, Stem Cell and Brain Research Institute U1208, Bron, France

## Abstract

The ability to attribute thoughts to others, also called theory of mind (TOM), has been extensively studied. Computationally, the basis of TOM in humans has been interpreted within the predictive coding framework and associated with activity in the temporo-parietal junction (TPJ). However, the evolutionary origins of these human mindreading abilities have been challenged since the concept was coined. Here we identify a brain region in the Rhesus macaque that shares computational properties with the human TPJ. We revealed, using a non-linguistic task and functional magnetic resonance imaging, that activity in a region of the macaque middle superior temporal cortex was specifically modulated by the predictability of social interactions. As in human TPJ, this region could be distinguished from other temporal regions involved in face processing. Our result suggests the existence of a precursor for the theory of mind ability in the last common ancestor of human and old-world monkeys.

---

The ability to attribute mental representations to others, called Theory of Mind (TOM^1^) is key to complex human social interactions^2,3^. TOM abilities in humans have been most notably associated with activity in the temporo-parietal junction (TPJ) and the medial prefrontal cortex (MPFC)^4,5^. The question of TOM’s evolutionary origins has, however, been disputed since the concept was first proposed^1,6,7^. This is partly due to the reliance of human TOM studies on linguistic stimuli^4^. But even when innovative non-verbal designs are employed, the interpretation of performances on TOM-like tasks across primate species are highly debated^2,7–9^.

Results from comparative anatomy studies suggest continuity rather than discontinuity in the anatomical organization of the primate social brain. For instance, the medial prefrontal cortex (MPFC) maintains a broadly similar organization in macaques and humans^10^, and human TPJ has a similar connectivity pattern to macaque middle Superior Temporal Sulcus (midSTS)^11^. Despite evidence that macaque midSTS and MPFC respond to social stimuli^12–19^, it remains unclear whether these regions support functions resembling TOM.

Theoretical developments in computational neuroscience suggest alternative ways to compare human and animal social abilities. Rather than looking for TOM itself in other species it may be profitable to seek evidence of more basic computational processes linked to TOM^20–22^. Computational models describe human TPJ and MPFC activation during social tasks within a predictive coding framework^3,23^. This framework predicts that deviations from expected social behaviors should lead to stronger activation in these areas.

## Macaques’ midSTS is modulated by social expectation

To investigate whether macaque brain areas signal deviation from social expectation, we presented 14 Rhesus macaques with a free-viewing functional Magnetic Resonance Imaging (fMRI) paradigm consisting of videoclips of macaques interacting socially. This approach has been successfully used to identify brain networks supporting social cognition in macaques ^24^ but has not yet been used to identify computations supported in those circuits. In our videos, social interactions either followed an expected scenario (e.g. continuous grooming or playing, Video 1-2) or were interrupted by an unexpected event (e.g. grooming or playing interrupted by a fight; Video 3-4). Several brain areas showed higher activation for the unexpected than expected social events, including regions belonging to the visual cortex and oculomotor-related regions (Supplementary Figure 1, Supplementary Table 1). Two clusters in the midSTS were also identified, which we will refer to as caudal midSTS and rostral midSTS (Fig. 1a, Supplementary Table 1). The rostral midSTS has often been associated with the macaque social brain^11,17,25^.

**Fig. 1.**
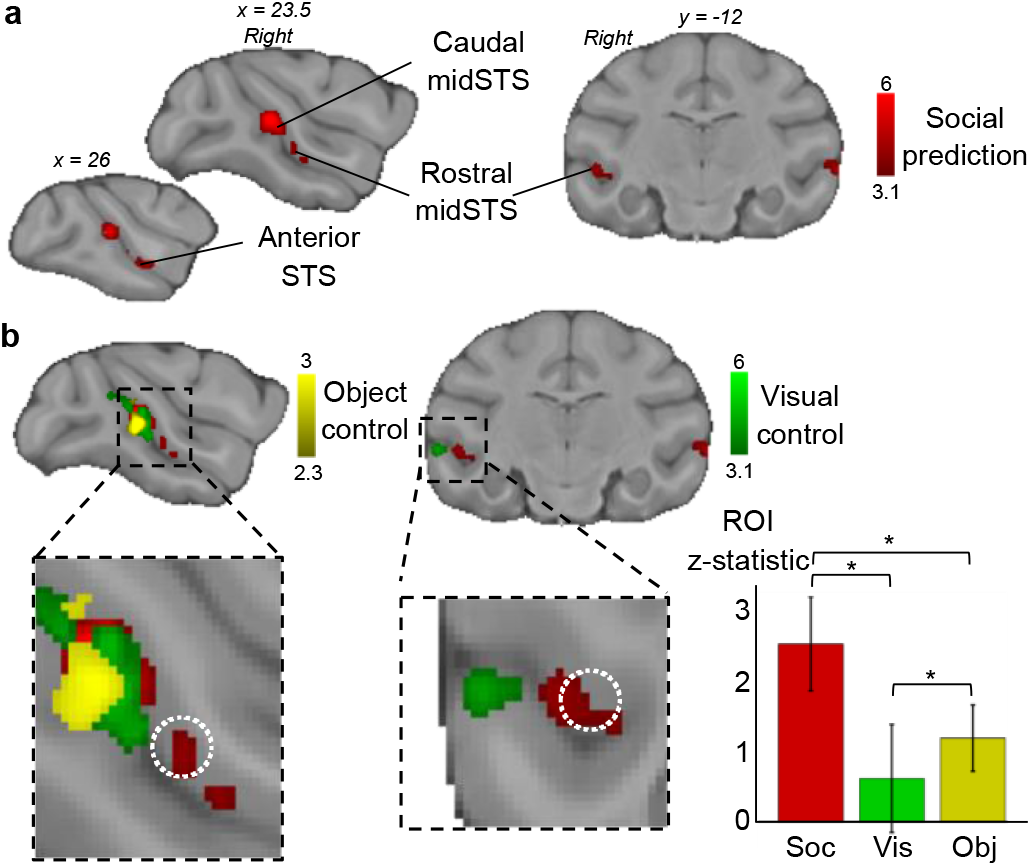
Modulation of macaque STS activity. **a**, Social prediction: group contrast of unexpected versus expected social interaction revealed activity in rostral and caudal midSTS (n=14, cluster-corrected at z>3.1, p<0.05 FWE corrected). **b**, Overlap between responses to social prediction and control conditions (visual control: n=14, cluster-corrected at z>3.1; object control: n=7, cluster-corrected at z>2.3 and both p<0.05 FWE corrected). The white dotted circle represents a macaque TPJ-like region identified previously^11^. Mean Z-statistic obtained in the ROI (white circle) for social prediction (soc), visual control (vis), object control (obj). Error bars represent standard deviation (Wilcoxon signed-rank test, Bonferroni corrected for multiple comparison p<0.05, social × visual: p=6.10^-43^, social × object: p=3.10^-43^, object × visual: p<1.10^-21^).

To rule out explanations in terms of basic visual features, we first contrasted the neural response to scrambled videos of unexpected versus expected social interactions which were matched in terms of luminosity and movement to the original videos (visual control). The visual control contrast elicited higher activation in the caudal midSTS but not in the more rostral part of the midSTS (Fig. 2b, Supplementary Table 1, Supplementary Fig. 2a). Unexpected social interaction videos contain by definition more unexpected movement and therefore it is not surprising that this visual control would recruit regions in caudal midSTS that include the motion-sensitive areas MT, FST and MST^25,26^.

**Fig. 2.**
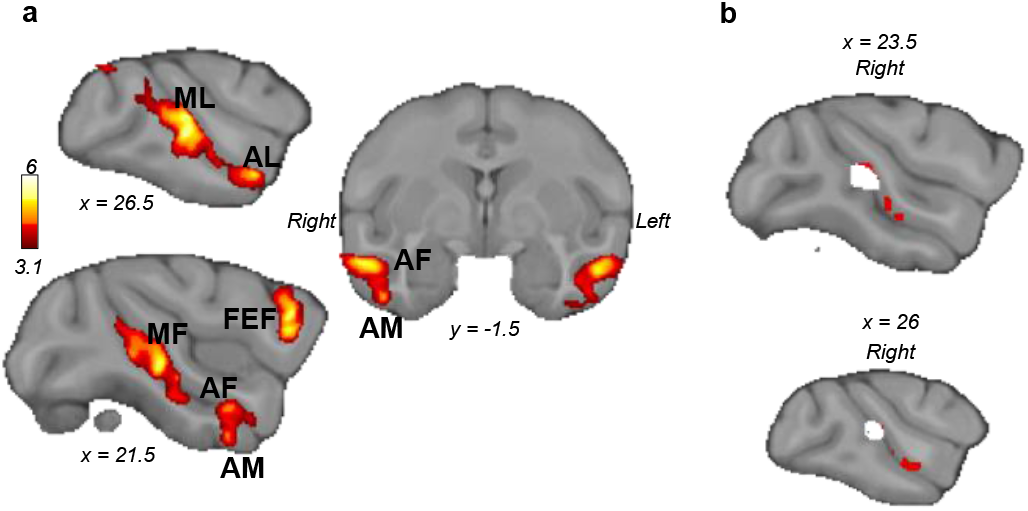
Face-responsive areas in macaques. **a,** Macaque group contrast of face versus scrambled pictures (n=14, cluster-corrected at z>3.1, p<0.05 FWE corrected). **b,** Conjunction analysis (white) of social prediction contrast activation (red) and face patches (cluster-corrected at z>3.1, p<0.05 FWE corrected).

We then tested the social specificity of the activation observed for social prediction in a subset of subjects (n=7/14, object control) using non-social scenes containing inanimate objects. To match closely with the social interaction videos, these videos were designed to represent situations with or without a departure from an expected and established physical regularity, such as the location, identity or movement. Regardless of whether we examined activity at the original threshold (z>3.1) or at a more liberal threshold (z>2.3) to account for the smaller number of animals, there was no evidence for activity in rostral midSTS but only in caudal midSTS for this object control (Fig. 1b, Supplementary Table 1, Supplementary Fig. 2b). A conjunction analysis between the social prediction contrast and each of our two control conditions (Supplementary Fig. 2c) confirmed the specificity of the modulation of activity by social predictability in rostral midSTS cluster.

From here on, we will refer to this specific rostral midSTS region as ‘social prediction area’ (SPA). It overlaps with cytoarchitectonically defined temporo-parieto-occipital association area (TPO) and PG associated area of STS (PGa)^27^. From this location, we can also rule out an overlap with body responsive areas which have been identified either posteriorly or ventrally to the SPA^24,28^. It has also recently been shown that strategic social signaling in the rostral midSTS involves a different set of neurons than the ones responding to faces and bodies^18^. Importantly, the rostral midSTS we identified corresponds to a midSTS region previously identified for its connectivity pattern most resembling that of human TPJ ^11^. Using this independently defined region of interest (ROI), we observed that social prediction induced significantly higher activation than control conditions (Fig. 1b).

To confirm that the social prediction modulation in the SPA was not due to a thresholding effect and illustrate the specificity of its activity, we performed the three contrasts (social prediction, visual control, object control) using the same independent ROI identified previously^11^ to restrict the statistics. We observed a significant activation in the ROI only in the social prediction contrast and not the two others with a cluster correction (Supplementary Fig. 3, top). Because the extent of this ROI is quite small, we also performed voxel correction which showed again the specificity of activation in this region for the social prediction contrast (181 voxels significant out of the 257 voxels of the ROI) and only a few voxels for the other two on the posterior edge of the ROI (12 for the visual control and 3 for the object control, Supplementary Fig. 3, bottom).

While we observed midSTS clusters bilaterally, some hemispheric differences were noticeable. The right caudal midSTS cluster, unlike the left caudal mid-STS, extended toward the end of the STS, including V4t on its ventral bank^27^. On the left hemisphere, the rostral midSTS cluster was located in a different area than the right SPA and had a more lateral position, extending from the dorsal bank of the STS to area TE on the lateral surface. To investigate whether the lack of social prediction modulation in the left SPA was indicative of a thresholding issue or a lateralized function, we defined a large ROI encompassing the whole STS around the coordinates of the previously mentioned midSTS region sharing neuroanatomical similarities with the human TPJ^11^. With the same social prediction contrast but restricted on either the left or right hemisphere of this enlarged ROI, we found that a cluster survives the statistical correction in both hemispheres. Rather than a purely lateralized function, these results show that the modulation by social prediction in the SPA was bilateral but less robust in the left hemisphere (Supplementary Fig. 4).

Finally, we conducted a separate free-viewing experiment with a different set of four monkeys. We collected sessions under different experimental conditions to verify that the social prediction modulation in SPA is robust to replication and to disruption of other brain regions. Due to the passive nature of the task, it was not possible to causally address the role of the midSTS. Instead we used repetitive Transcranial Ultrasound Stimulation (TUS) protocol to disturb brain activity over key regions of interests to test for potential confounding effects. Repetitive TUS could disrupt the normal activity of the stimulated region for at least two hours post-stimulation^29^. With the sessions with no-prior stimulation, we replicated the previous results, revealing the same rostral midSTS region as specifically modulated by social prediction (Supplementary Fig. 5, Supplementary Table 2). In the replication, the visual and object control contrasts did not yield any significant results in rostral mid-STS and there was no conjunction with the social prediction contrast (Supplementary Fig. 6).

In both the original and replicated studies, we observed a cluster just anterior to the genu of the arcuate sulcus, an oculomotor region often referred to as the Frontal Eye Field (FEF). To rule out a putative attentional or oculomotor confound with the social prediction modulation, we used, prior to the awake fMRI data acquisition, a repetitive TUS stimulation of the FEF. In separate sessions the Anterior Cingulate Cortex (ACC), a region known for its role in social cognition^12,17,24^, was targeted as an active control region. The stimulations do perturb some brain network as they have an effect on two relevant contrasts: a simple visual contrast (videos versus black screen) and a social contrast (social videos versus scrambled) (Supplementary Fig. 7). However, in our contrast of interest, the social prediction, no difference between stimulation and non-stimulation sessions could be observed. These results show that SPA was modulated by the predictability of social interaction, independently of attentional or oculomotor effect led by the FEF. They confirmed the social specificity of the activity modulation in the SPA.

## Relationship of SPA with the face responsive brain network

To further test specificity of SPA responses and their relationship with known STS functions, we investigated how SPA are related to face patches, a set of face-responsive areas located in STS and inferotemporal cortex ^30^. We analyzed awake fMRI data from a face localizer collected in our initial group of 14 Rhesus macaques. Our localizer consisted of pictures of neutral and emotional (e.g. lip-smacking, open mouth...) macaque faces and their scrambled equivalent during fMRI. This method has been shown to identify the face-responsive brain regions as opposed to the face-selective brain regions by using a localizer combining face, body and object pictures ^31^. In 12 out of 14 animals we were able to identify all six face patches previously reported^19,30^: Posterior Lateral (PL), Middle Lateral (ML), Middle Fundus (MF), Anterior Lateral (AL), Anterior Fundus (AF) and Anterior Ventral (AM) (Fig. 2a, Supplementary Fig. 7, Supplementary Table 3).

A conjunction analysis revealed no significant overlap between face patches and SPA (Fig. 2b). At the single-subject level, we noticed SPA peaks tended to be located in a more dorsal/fundus section of mid-STS, and therefore in a distinct cytoarchitectonic area compared to face patches (Supplementary Fig. 8). Our results are supported by recent findings showing that neurons in the ventral bank of the midSTS signal selectively cooperative social behavior, independently from visual sensitivity to faces and bodies^18^.

We then conducted a resting state fMRI analysis to determine the relationship between the SPA and the face patches. We computed the functional connectivity profiles of macaques’ SPA with both full correlation as available in humans and a more specific partial correlation. The full correlation revealed that macaques’ SPA coupling with face responsive regions and other visual areas (Fig. 3a) which was absent for human TPJ connectivity profile (coordinates from^11^ HCP resting-state data^32^, Fig. 3b). However, computing the partial connectivity, by regressing out the time series of all face patches, reveals that SPA is specifically coupled with dorsal STS, posterior cingulate and prefrontal cortex, resembling the human TPJ connectivity profile (Fig. 3c). Similarly, computing the partial connectivity of the face patches, by regressing out the time series of the SPA (and its anterior section) revealed a network involving mostly STS and the visual cortex (Fig. 3d). In summary, connectivity results provide further evidence for the distinction of face patch and SPA systems, but also reveal stronger interactions between the two systems in macaques than in humans.

**Fig. 3.**
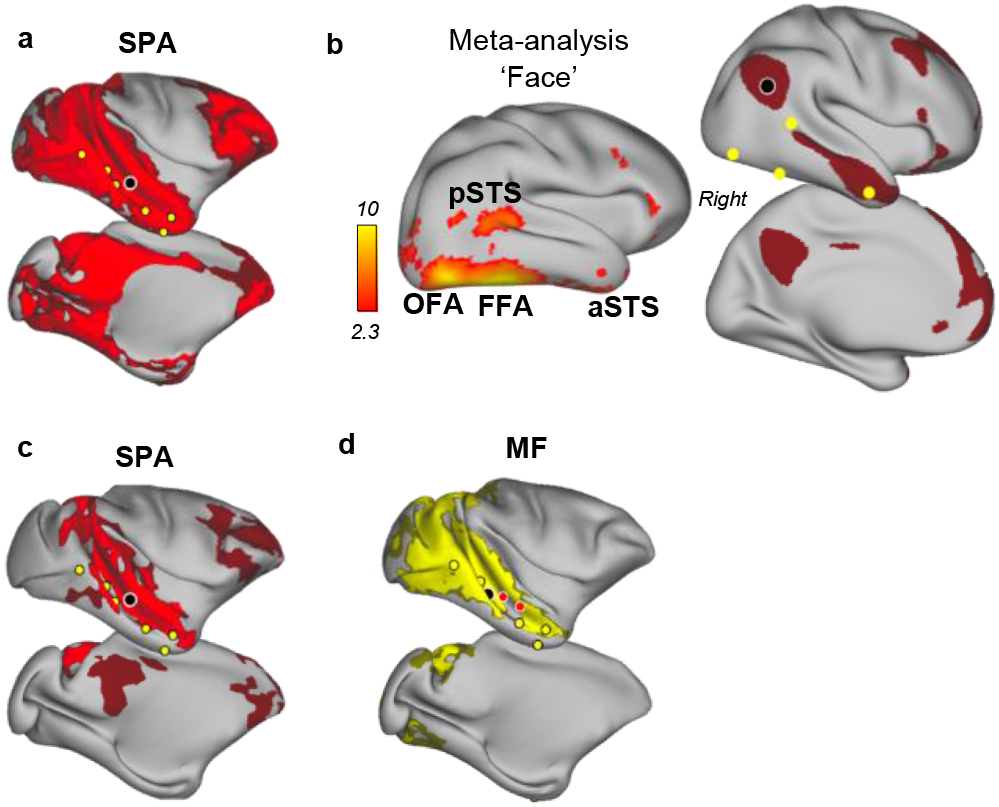
Face-patch system and resting-state functional connectivity in macaques and humans. **a,** Resting-state connectivity associated with SPA (black circle) from a full correlation to the whole brain (face patches are yellow circles). **b,** Human comparison. Right: meta-analysis results (Neurosynth) for ‘face’, displayed on right hemisphere (pSTS: posterior STS, OFA: Occipital Face Area, FFA: Fusiform Face Area). Left: resting-state connectivity of TPJ (Cohen’s d effect size thresholded at 0.6). **c,** Resting-state connectivity associated with SPA (black circle) from a partial correlation to the whole brain while accounting for face patches connectivity. **d,** Resting-state connectivity associated with MF (black circle) from a partial correlation to the whole brain while accounting for SPA connectivity (SPA and its anterior section are red circles). For all macaque resting state: n=12, TFCE corrected, FWE corrected at p<0.01 in bright color and 0.01<p<0.05 in dark color.

## Discussion

Overall, our results revealed a brain region in macaques’ rostral midSTS that is specifically sensitive to expectation violation during free-viewing of social scenes. This region is distinct from previously identified functional module in the STS called face, gaze-following, or body patches^19,28,30,33^. Its location on the dorsal bank / fundus of the STS is compatible with a functional module identified as being responsive to natural social scenes^34^. Here, we were able to characterize a computational property associated with this region. We interpret this response in a predictive coding framework providing the signature of the neuro-computational mechanism supporting mentalizing abilities in humans^3^. Evidence for this type of coding has been uncovered in adjacent regions of the temporal cortex for processing non-social information in macaques^35^. Furthermore, the midSTS region sensitive to prediction in the social domain corresponds to the region that was previously shown to share similar connectivity profiles with the human TPJ^11^.

Our approach, built on theoretical debates about cross-species differences in TOM^2,7,21^, provides evidence for the existence in the last common ancestor to humans and macaques of a precursor neural architecture supporting computations that have been associated with TOM in human TPJ^21^. Unlike in human studies^4,5^, our social prediction analysis did not reveal any change of activity in macaque MPFC. This may reflect the nature of the passive free-viewing tasks compared to the active decision-making tasks used in humans^23,36^.

Our results suggest an evolutionary trajectory in brain organization that in humans has resulted in area TPJ. The connectivity of face-responsive areas and the SPA differs in both humans and rhesus macaques but the two circuits are more integrated in macaques; macaque SPA retains connectivity to face patches while human TPJ shares little connectivity with the face-responsive system. These between-species differences might reflect greater specialization of social brain areas such as TPJ in humans that may have occurred in association with the expansion and reorganization^37^ of the temporal cortex since the last common ancestor to humans and old-world monkeys 25 to 29 millions years ago.

## Methods

### Data acquisition

#### Animals

14 healthy Rhesus macaque monkeys (*macaca mulatta*, 13 males, 1 female) performed a set of free-watching tasks over a period of six months. All procedures were conducted under licenses from the United Kingdom (UK) Home Office in accordance with the UK Animals (Scientific Procedures) Act 1986. See Supplementary Table 4 for a detailed account of the number of runs per conditions and per monkeys.

#### Stimuli

Pictures and videos recorded at the breeding center and at the Oxford research colony were the basis of the video clips used in four experimental conditions. In addition, two other experimental conditions based on non-social stimuli were also used. Altogether these six conditions and an awake resting-state acquisition (not included in this study) were presented in pseudo-randomized order. Three conditions described below have been used for the purpose of the current study. No more than three repetitions of a given condition was presented per day, the same condition was never repeated consecutively, and two different orders of presentation of the videos/pictures for a given condition were used to further limit habituation. For all conditions, the animals were not asked to fixate their gaze, to conserve the most natural behavior. No reward delivery occurred during the presentation of stimuli. Reward was instead delivered in between two runs to maintain animal attention to the stimuli.

First, we selected videos containing expected (ex: grooming or playing) and unexpected (ex: unexpected deviation from grooming or playing) social behaviors which were highly ecologically valid for the monkeys. The videos were presented for 5.5 seconds each and were combined in a 12-second block with 0.5 second before each video. Each block was followed by 10 seconds of rest (black screen). We presented three blocks of social unpredicted, three blocks of social predicted and three blocks for each of their scrambled versions respectively in a random order.

Based on a similar principle (deviation from expected interaction) we created videos showing expected and unexpected object interactions based on simple physical regularity. In keeping with the social videos, object scenes showed events that could be unpredicted based on either location (object appearing at unexpected location), identity (a new object appears), or movement (sudden change in movement patterns shown up to now). For instance, a video in which objects are falling at constant rate is considered predictable while an unpredictable scenario would see this rate suddenly changed without an obvious cause. The timings for these conditions were the same than for the social prediction. We presented three blocks of object unpredicted interactions, three blocks of object predicted interactions and three blocks for each of their scrambled version respectively. This task was only done on seven of the 14 monkeys.

For the face localizer, the task followed a block design with each block of 12 seconds consisting of the presentation of eight images for 1 second each followed by 500 ms of black screen. A resting period of 10 seconds (black screen) was inserted between the face blocks. Each run was composed of three blocks of neutral faces, three blocks of emotional faces (aggressive or lip-smacking) and six blocks of scrambled faces. This type of face localizer is known to capture face-responsive areas ^31^.

#### Awake and anaesthetized fMRI

The fMRI data were acquired in a horizontal 3 Tesla MRI scanner with a full-size bore using a four-channel, phased-array, receive-only radio-frequency coil in conjunction with a local transmission coil (Windmiller Kolster Inc, Fresno, USA). The animals were head-fixed in a sphinx position in an MRI-compatible chair (Rogue Research, CA). fMRI data were acquired using a gradient-echo T2* echo planar imaging (EPI) sequence with the following parameters: 1.5 × 1.5 × 1.5 mm resolution, 36 axial interleaved slices with no gap, TR of 2280 ms, TE of 30 ms and 130 volumes per run. Proton-density-weighted images using a gradient-refocused echo (GRE) sequence (TR = 10 ms, TE = 2.52 ms) were acquired as reference for offline image reconstruction.

Resting-state fMRI data and anatomical scans were collected under anesthesia for the same animals according to a previously used protocol^10^. fMRI Resting-state connectivity patterns are well conserved under anesthesia^38^, and have been used for conducting human-macaque comparisons^10,11,38^. Anesthesia was induced using intramuscular injection of ketamine (10 mg/kg) combined with either xylazine (0.125–0.25 mg/kg) or midazolam (0.1 mg/kg) and buprenorphine (0.01 mg/kg). Macaques also received injections of atropine (0.05 mg/kg), meloxicam (0.2 mg/kg), and ranitidine (0.05mg/kg). Anesthesia was maintained with isoflurane. Isoflurane was selected because it has been demonstrated that resting-state networks are still present using this agent for anesthesia ^38^. The anesthetized animals were placed in an MRI-compatible stereotactic frame (Crist Instrument) in a sphinx position within a horizontal 3T MRI scanner with a full-size bore. The same coils as for awake scans were used for data acquisition. Whole-brain BOLD fMRI data were collected using the following parameters: 1.5 × 1.5 × 1.5 mm resolution, TR of 2280 ms, TE of 30 ms, 36 axial interleaved slices with no gap and 1600 volumes. Structural scans were acquired in the same session using a T1-weighted MP-rage sequence (no slice gap, 0.5 × 0.5 × 0.5 mm resolution, TR of 2500 ms, TE of 4.01 ms and 128 slices).

### Preprocessing

All data were preprocessed and analyzed using tools from the FMRIB Software Library (FSL, version 5.0.10) ^39^, the Advanced Normalization Tools (ANTs, version 2.1.0) and the Connectome workbench software (www.humanconnectome.org). We also used MATLAB (version R2016a, The MathWorks, Inc., Natick, Massachusetts, United States) and bash codes from the Magnetic Resonance Comparative Anatomy toolbox (MrCat, www.neuroecologylab.org) and custom-made codes.

#### Task-fMRI preprocessing

Task-fMRI data were preprocessed following a dedicated non-human primate fMRI processing pipeline as part of the MrCat toolbox. In short, after offline SENSE reconstruction of the EPI image (Windmiller Kolster Scientific, USA), motion-induced time-varying slice distortions were corrected using restricted non-linear registration, first to a run-specific high-fidelity EPI, then to each animal’s T1w structural image, and finally to group-specific template in CARET macaque F99 space ^40^. Brain extraction, bias-correction, and template registration of the T1w structural image were achieved in an interdependent iterative approach. The resultant high-fidelity removal of non-brain tissue could be back-projected to the EPI following non-linear registration. A nuisance regressor design matrix was created to account for volumes with excessive movement, signal variability associated with motion-induced distortion artifacts and non-brain noise components. For the video tasks, we did not use the regressors for the non-brain component as they were correlated with the timing of the task. Further steps were implemented using the FEAT toolbox. We performed spatial smoothing using a Gaussian of 3 mm FWHM (full-width at half minimum) kernel, grand mean intensity normalization and high-pass temporal filtering (Gaussian-weighted least-squares straight-line fitting, with sigma = 100 s).

#### Resting-state fMRI preprocessing

The detailed preprocessing pipeline for the resting-state fMRI has been described elsewhere ^29,41^. Briefly, after reorientation to the same convention for all functional EPI datasets, the first volumes were discarded to ensure a steady radio frequency excitation state. EPI timeseries were motion corrected using MCFLIRT ^42^. Brain extraction, bias-correction, and registration were achieved for the functional EPI datasets in an interdependent iterative manner. The mean of each functional dataset was registered to its corresponding T1w image using rigid-body boundary-based registration (FLIRT ^42,43^). EPI signal noise was reduced both in the frequency and temporal domain. The functional timeseries were high-pass filtered with a frequency cut-off at 2000 s. Temporally cyclical noise, for example originating from the respiration apparatus, was removed using band-stop filters set dynamically to noise peaks in the frequency domain of the first three principal components of the timeseries. To account for remaining global signal confounds we considered the signal timeseries in white matter and meningeal compartments, and their confound parameters were regressed out of the BOLD signal for each voxel. Following this confound cleaning step, the timeseries were low-pass filtered with a cut-off at 10 s. The data were transformed to the surface space using the F99 template and spatially smoothed using a 2.8 mm FWHM gaussian kernel, while considering the folding of the cortex. Lastly, the data timeseries were demeaned to prepare for functional connectivity analyses.

### Analysis

#### Contrasts

For the awake fMRI, the first-level analysis was carried out using FEAT for each run ^44,45^. Simple generalized linear model (GLM) designs were defined. For the social prediction task, we used four Explanatory Variables (EVs), accounting for the social expected scene, social unexpected scene, and one for each of their scrambled versions. The main contrast of interest was between social unpredicted versus social predicted. We defined one more contrast as the scrambled unpredicted versus scrambled predicted to control for activity related to visual features (e.g. motion, luminance). We used a similar approach for the object prediction. For the face task, four EVs were used to account respectively for the neutral face blocks, the emotional face blocks, the neutral scrambled blocks and the emotional scrambled blocks. The main contrasts were defined as face images versus scrambled images and emotional faces versus neutral faces.

In each task on top of the main contrasts, we defined a control contrast to detect neural activation when an image or video was present on the screen compared to rest period, to confirm whether the monkeys were engaged during the task. Indeed, as the task did not provide reward to the animals, they could disengage and fall asleep. We therefore excluded runs in which this control contrast elicited no or limited activation in the visual cortex. This method excluded 5 runs for the social prediction task and 3 runs for the object prediction task.

We applied a gamma hemodynamic response function convolution with a phase of 0 seconds, a standard deviation of 1.5 seconds and a delay of 3 seconds and the same temporal filtering as for the data. The movement regressors previously described were also used as additional confounds.

In the second-level analysis, after registration to standard space we pooled together runs from the same monkeys. A fixed-effect analysis was performed at the subject level. Finally, a third-level analysis was carried out to obtain the results at the group level using FLAME 1 as mixed-effects analysis with a cluster-forming z-threshold of 3.1 and corrected for Family-Wise Error (FWE) at p<0.05. The z-thresholds were chosen according to previous literature ^46^ which advise to use the threshold of 3.1 with Flame 1 mixed-effect to avoid false positives. To test for a potential overlap of object prediction with social prediction, we used a more liberal threshold at z=2.3. In fact, when no complete overlap is expected, as here, this approach increases the sensitivity of the test allowing more stringent inferences.

#### Conjunction

We verified the specificity of the modulation by the social prediction videos by performing a series of conjunction analysis at the group level. All conjunctions are performed according to previous literature ^47^. We defined an STS mask comprising the grey matter of the STS excluding the very posterior parietal portion, to restrict the conjunction and set the cluster forming threshold at z=3.1 and p<0.05. For the conjunction between object prediction and social prediction we used only the same seven animals available in both datasets. Because no significant conjunction was found between the object and social prediction at the z>3.1 threshold, we lowered the threshold to 2.3, as above to increase the sensitivity and account for the smaller number of animals in this condition.

#### Comparison of mean uncorrected z-statistic

To further confirm that this result was not due to a thresholding effect, we conducted additional analyses. We defined a Region of Interest (ROI) around the coordinates found in an anterior study ^11^ (most similar connectivity profile to human TPJ) with a 5 voxel radius. First, we computed the mean uncorrected z-statistic across voxels in this ROI for our three conditions (social prediction, visual control and object control). The standard deviation is defined as the square root of the variance of the z-statistic. We performed a Wilcoxon signed-rank test between conditions and corrected for multiple comparison using the Bonferroni method. Secondly, we performed the same third-level contrasts as before but restricting the statistics to the rostral midSTS ROI as defined before. Because the extent of this ROI is quite small, we performed both cluster- and voxel-thresholding corrections.

#### Hemispheric and regional specificity

We also investigated the hemispheric specificity of the social prediction modulation by analyzing the same contrast with an ROI either on the left or on the right hemisphere as performed in the literature ^48^. The ROI was defined as a coronal mask (5 slices) encompassing the whole STS at the level of the small ROI mentioned earlier, around the coordinates found in the anterior study ^11^. This ROI was defined to overcome the issue of thresholding by reducing the number of voxels and to enlarge the search area so that we could capture clusters even if they were overlapping the borders of the small ROI (accounting for inter-individual differences).

The MPFC has also been identified as part of the social brain in macaques^17^. Therefore, we conducted another ROI analysis targeting the Anterior Cingulate Cortex (ACC) to restrict the statistics to this previously identified region^17^. No activity modulation of the ACC by the social prediction was revealed with this analysis.

#### Resting-data fMRI analysis

For the anaesthetized resting-data fMRI, in each monkey individually, we identified bilateral face patches from peak activation at the second-level analysis and based on the definitions of a previous study ^30^. We obtained the middle fundus (MF) and middle lateral (ML) in all monkeys, the anterior lateral (AL), the anterior fundus (AF) and the anterior medial (AM) in 13 monkeys and the posterior lateral (PL) in 12 monkeys. When the face patch was present on only one hemisphere we defined the opposite hemisphere face patch as its symmetric voxels. We carried on the analysis on the 12 monkeys where we could find all the face patches in at least one hemisphere. Each face patch location was mapped to surface space and a ROI was made of a circle of 2mm geodesic distance giving all ROIs the same size. We followed the same procedure for the social prediction area (SPA) and defined an anterior SPA ROI which was part of the same cluster but could be found in all monkeys, insuring that we cover the entirety of the modulation location. We extracted the time series of each of these ROIs (six for face patches, two for social prediction) and computed their correlation with timeseries of the whole brain. We also performed a partial correlation where we regressed out the mean time series of all face patches from the SPA and the time series of the SPA from the face patches to obtain their specific connectivity. We then computed the correlation of these more specific time series to the whole brain. We therefore obtained two maps describing how each ROI connects to the rest of the brain for each monkey using both full correlation and partial correlation. We merged all monkeys for each seed and performed a non-parametric permutation inference using PALM ^49^ and performing the maximum number of permutations (in this case sign-flipping for a one-sample t-test). Clusters were defined with the threshold-free cluster enhancement (TFCE) method which enhances the cluster-like structures but keep the voxel dimension of the data and were corrected for multiple comparison using the Famil-Wise-Error method.

For visualization, some of the results were projected onto the F99 surface using tools from the HCP workbench and the inflated surfaces from a published study ^50^ (Supplementary Fig 8).

### Human Data

For the face task, we used the Neurosynth platform (Created and maintained by Tal Yarkoni, supported by NIH award R01MH096906) for automated meta-analysis that we probed with the word ‘faces’. The resulting meta-analysis map from 864 studies was then z-stats thresholded at 2.3 and projected onto a standard MNI surface. The map is corrected using a false discovery rate (FDR) approach, with an expected FDR of 0.01.

For the resting-state human study, data were provided by the Human Connectome Project, WU-Minn Consortium (Principal Investigators: David Van Essen and Kamil Ugurbil; 1U54MH091657) funded by the 16 NIH Institutes and Centers that support the NIH Blueprint for Neuroscience Research; and by the McDonnell Center for Systems Neuroscience at Washington University. We specifically used the group-average structural and functional MRI data from the HCP S1200 data release (March 2017). This dataset, available on-line at https://www.humanconnectome.org, allowed us to access task-related data but also resting-state connectivity network and atlases. The connectivity of TPJ was obtained from a ROI of 2mm geodesic distance around the TPJ coordinates defined as a in previous study ^51^.

### Replication and Transcranial Ultrasound Stimulation

One year after the first acquisition batch, we were able to acquire additional data for four animals (T2, T3, T4 and an additional monkey V1). Therefore, we conducted a replication study using 6 sessions for each of the conditions per animal (social prediction: 24 sessions, visual control: 24 sessions, object control: 24 sessions). We followed the exact same procedure, except for some technical acquisition and analysis details that we describe here. Data were collected with a 3-Tesla MRI scanner with a full size bore and we used the four-channel, phased-array, receive-only radio-frequency coil in conjunction with a local transmission coil (Windmiller Kolster Inc, Fresno, USA). We used the exact same acquisition protocol. Concerning the analysis, we restrained our analysis to two levels, because of the limited amount of data and because this is the most commonly used approach when having the same number of sessions for each animal. At threshold level 3.1, we did not obtain any significant result, but this was expected considering the lower amount of data. Therefore, we lowered the threshold to 2.3 and performed the same conjunction analysis and calculated the mean uncorrected z-statistic across voxels in this ROI as in the initial study.

To assess if attentional or oculomotor related neural activity could explain the modulation by social prediction in the SPA, we performed Transcranial Ultrasound Stimulation (TUS) on the same macaques used for the replication just prior to the fMRI free-viewing task. We stimulated the Frontal Eye Field (FEF) as it is involved in attention and oculomotor movement such as saccades^52,53^, and was also revealed in our social prediction analysis. As a control region, we stimulated the ACC which is involved in the extended social brain. The impact of TUS on FEF and ACC and their consequence on behavior have already been demonstrated^41,52–54^. We also collected control data for which no stimulation was performed (note that these are the data used in the replication). For these three stimulation conditions, we acquired 6 runs per monkeys per conditions (social prediction, visual control, object control). Control days were interleaved with TUS sonication days. TUS was performed using the same protocol as previously published ^54,55^ adapting the focal depth of the transducer to the desired coordinates. Note that one FEF session for one animal was conducted with a higher intensity (60% duty cycle instead of 30%) which resulted in a localized skin trauma. A sequential stimulation was performed to target the left and right FEF^55^. A unique stimulation was performed on the midline for achieving a bilateral ACC stimulation^54^. Briefly, a single-element ultrasound transducer was used for 40 s. It was positioned with the help of Brainsight neuronavigation system (Rogue Research) so that the focal spot was centered on the targeted brain region, namely the FEF on the anterior bank of the arcuate sulcus (left FEF MNI coordinates +/-SD: × =-14.4+/-0.9, y = 4.9+/-2.5, z = 13.3+/-1.4; right FEF: x=15+/-1.2, y = 4.2+/-1.6, z= 11.8+/-1.5) and the controlled region: the ACC rostral to the genu of the corpus callosum (MNI coordinates +/-SD : × = 0 +/- 0.9, y = 15.5 +/- 1.5, z = 6.5 +/- 1.0). fMRI data acquisition, preprocessing and analysis were performed as described for the replication. To compare control condition contrasts with stimulation condition contrasts we performed a Two-Sample Paired T-Test, regressing out the mean of each subject so that it would not interfere with the estimation of the difference between stimulation conditions. To assess that the stimulations had any effect, we compared a simple visual contrast (videos versus black screen) and a social contrast (social videos versus scrambled). Having established that stimulations did change some of the brain task-related modulation, we compared the contrast of interest: the social prediction. We used a whole brain analysis as well as an ROI analysis. This ROI combined the left and right ROI defined for the hemispheric analysis resulting in a coronal mask encompassing the whole STS bilaterally at the level of the small ROI mentioned earlier. This ROI was defined to overcome the issue of thresholding and inter-individual difference.

## Acknowledgments

We thank Lev Tankelevitch, Caroline I. Jahn, Alessandro Bongioanni and Davide Folloni for their important contribution to the development of awake fMRI preprocessing pipeline. Human Data were provided by the HCP, WU-Minn Consortium (Principal Investigators: D. Van Essen and K. Ugurbil; 1U54MH091657) funded by the 16 NIH institutes and centers that support the NIH Blueprint for Neuroscience Research and by the McDonnell Center for Systems Neuroscience at Washington University.

Funding: L.R. is supported by funding from the Biotechnology and Biological Sciences Research Council (BBSRC) [BB/M011224/1]; M.S. is supported by the Austrian Science Fund: Erwin Schroedinger Fellowship [FWF-J4009-B27]; L.V. is supported by a Marie Curie Intra-European Fellowship within the European Union’s 7th Framework Programme [MC-IEF-623513] and funding from the Wellcome Trust [WT100973AIA]; A.M. is supported by an MRC grant [MR/P024955/1] and the Wellcome Trust centre grant [203139/Z/16/z]; M.K. is supported by the Centre National de la Recherche Scientifique, Mission pour l’Interdisciplinarité; M.F.S.R is supported by MRC grant [G0902373]; R.B.M. is supported by the Biotechnology and Biological Sciences Research Council (BBSRC) UK [BB/N019814/1] and the Netherlands Organization for Scientific Research NWO [452-13-015]; J.S. is supported by a Sir Henry Dale Wellcome Trust Fellowship [105651/Z/14/Z], IDEXLYON “IMPULSION 2020 grant (IDEX/IMP/2020/14) a Hayward JRF from Oriel College, University of Oxford and the Labex CORTEX ANR-11-LABX-0042 of Université de Lyon; The Wellcome Centre for Integrative Neuroimaging is supported by core funding from the Wellcome Trust [203139/Z/16/Z] and Wellcome Strategic award [WT101092MA];

## Supplementary Data

**Supplementary Figure 1.**
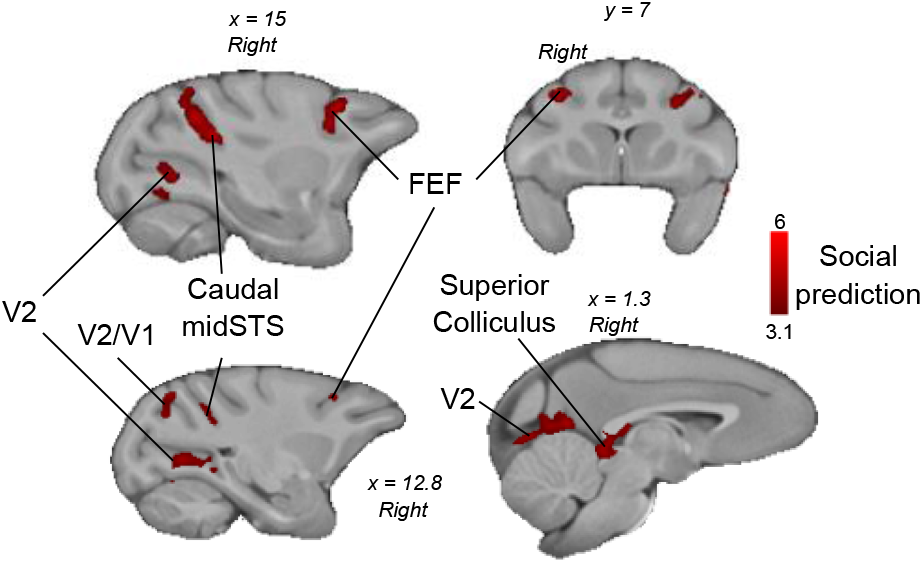
Social prediction contrast. Group contrast of unexpected versus expected social interaction also revealed activity in Frontal Eye Field (FEF), Superior Colliculus, Visual area V2 and Visual area V2/V1 (n=14, cluster-corrected at z>3.1, p<0.05 FWE corrected).

**Supplementary Figure 2.**
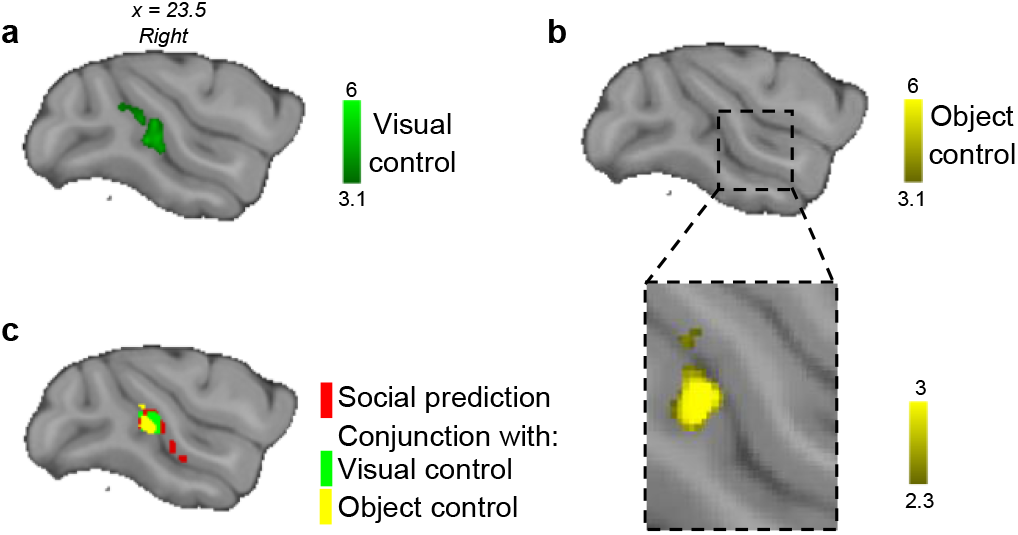
Control conditions. **a,** Visual control: group contrast of unexpected versus expected scrambled social scenes revealed activity in caudal midSTS only (n=14, cluster-corrected at z>3.1, p<0.05 FWE corrected). **b,** Non-social prediction (object) control: group contrast of unexpected versus expected object scenes revealed no activity (n=7, cluster-corrected at z>3.1 and p<0.05 FWE corrected). At lower threshold (insert), the contrast revealed activity in caudal midSTS only (cluster-corrected at z>2.3, p<0.05 FWE corrected). **c,** Conjunction results between the social prediction contrast and the control contrasts (cluster-corrected at z>3.1 for visual feature control and at z>2.3 for object control, p<0.05 FWE corrected).

**Supplementary Figure 3.**
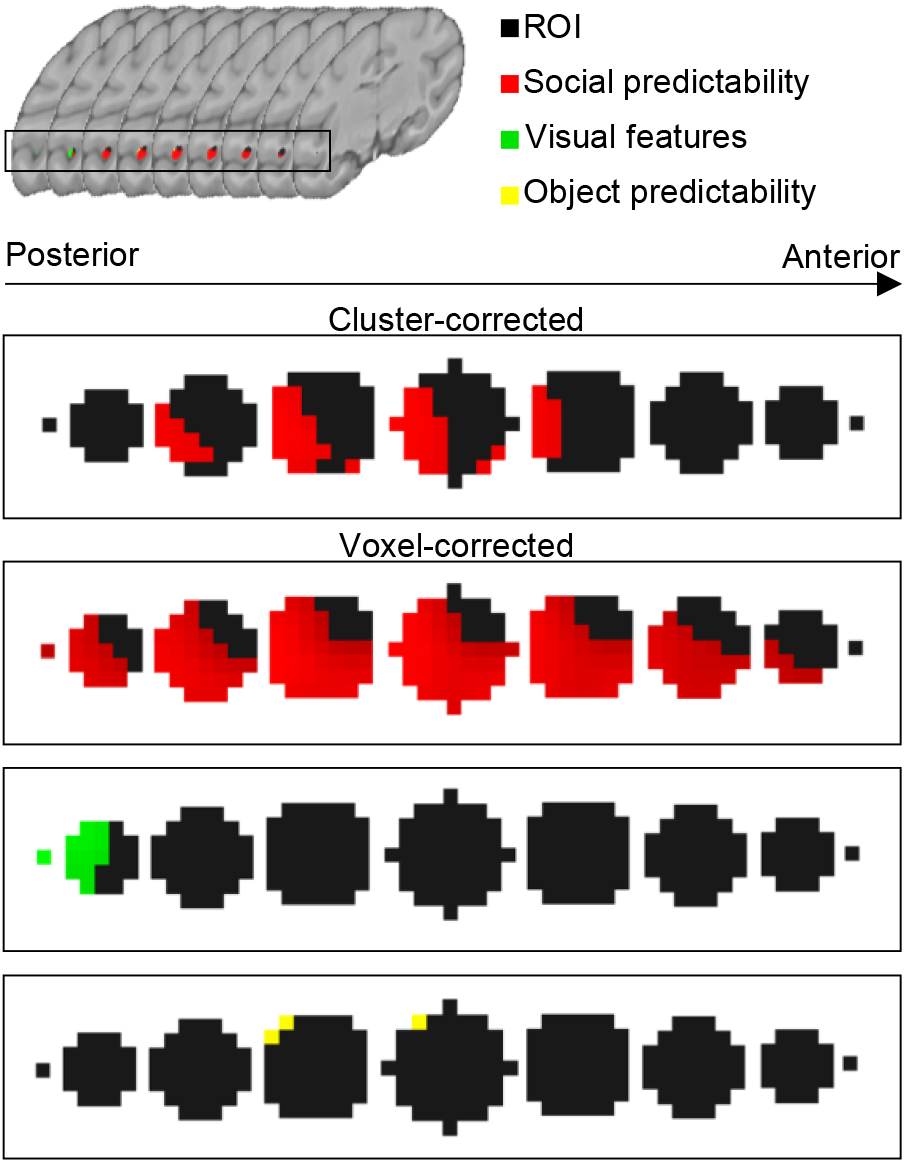
ROI analysis cluster and voxel corrected. Representation of the midSTS ROI from Mars et al (2013), from posterior to anterior coronal slices. When cluster-corrected (z>3.1) only the social prediction contrast was significant. When voxel-corrected (p<0.05), a few voxels in the two other controls were significant.

**Supplementary Figure 4.**
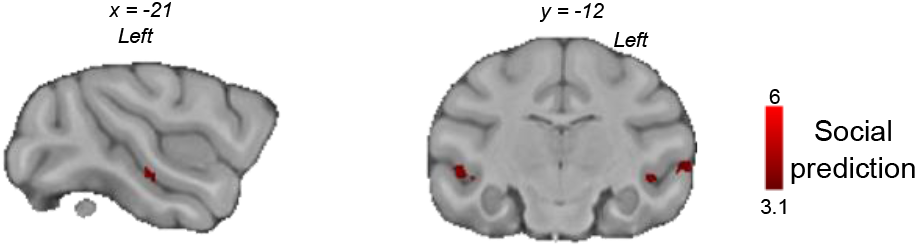
Hemispheric analysis. Social prediction: group contrast of unexpected versus expected social interaction restricted on a rostral midSTS ROI revealed activity in rostral on the left hemisphere midSTS (n=14, cluster-corrected at z>3.1, p<0.05 FWE corrected).

**Supplementary Figure 5.**
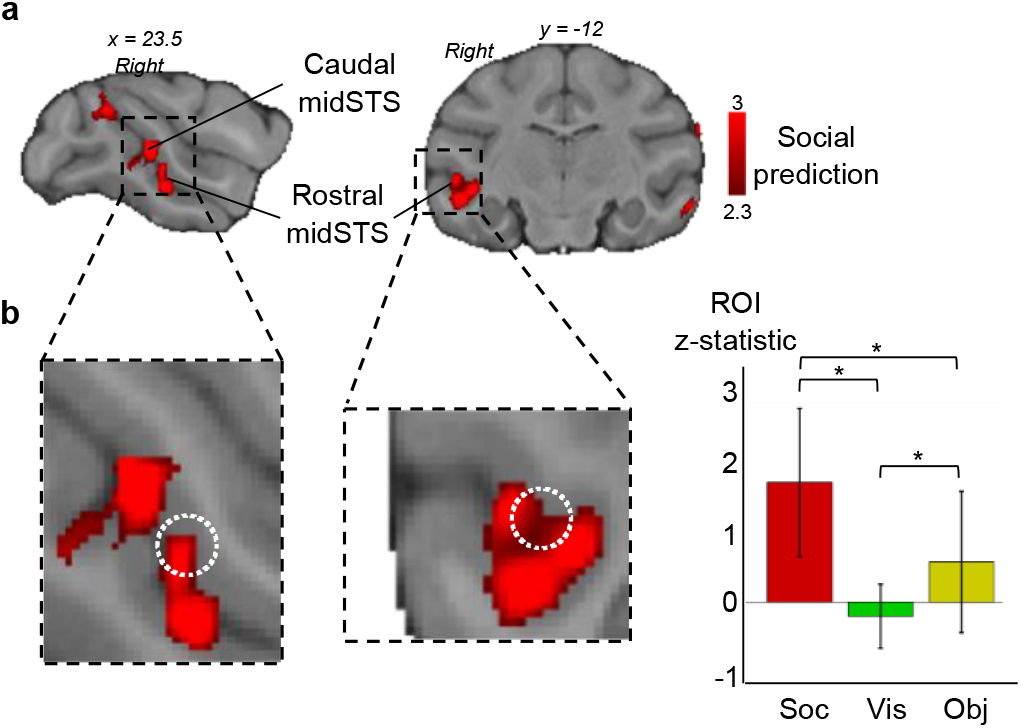
Replication of the modulation of macaque STS activity. **a**, Social prediction: group contrast of unexpected versus expected social interaction revealed activity in rostral and caudal midSTS (n=4, cluster-corrected at z>2.3, p<0.05 FWE corrected). **b**, Inserts show the white dotted circle representing a macaque TPJ-like region identified previously^11^. Mean Z-statistic obtained in the ROI (white circle) for social prediction (soc), visual control (vis), object control (obj). Error bars represent standard deviation (Wilcoxon signed-rank test, Bonferroni corrected for multiple comparison p<0.05).

**Supplementary Figure 6.**
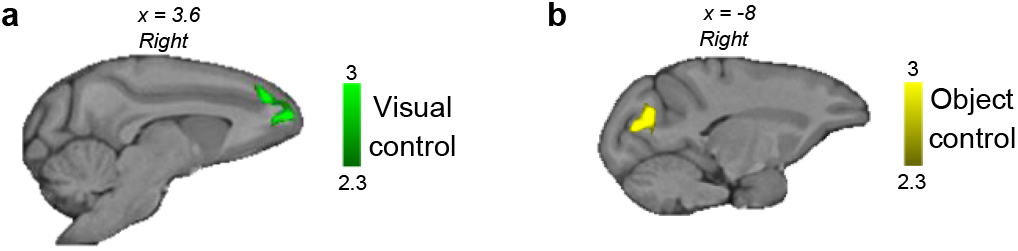
Replication of the control conditions. **a,** Visual control: group contrast of unexpected versus expected scrambled social (n=4, cluster-corrected at z>2.3, p<0.05 FWE corrected). **b,** Non-social prediction (object) control: group contrast of unexpected versus expected object scenes (n=4, cluster-corrected at z>2.3 and p<0.05 FWE corrected).

**Supplementary Figure 7.**
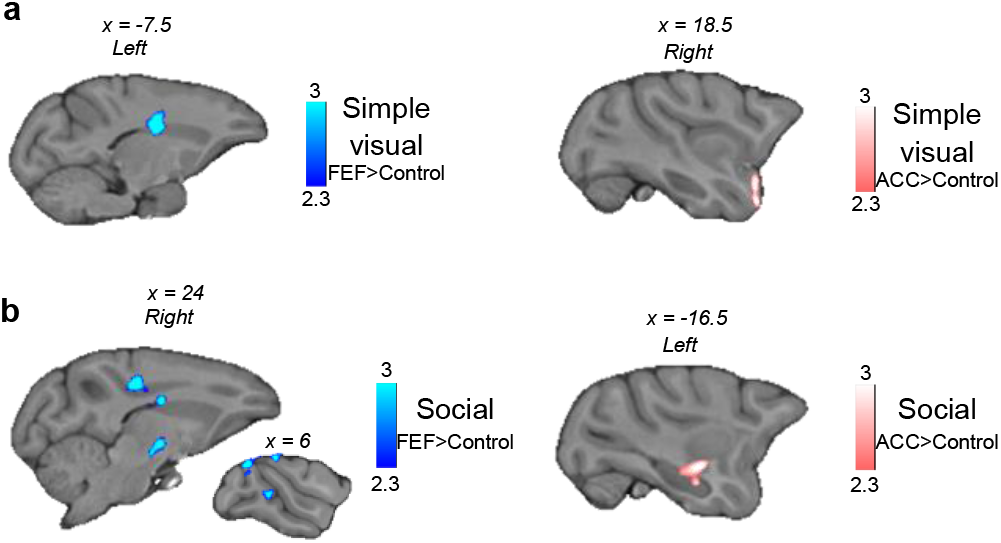
Effect of ultrasound stimulation. **a,** Simple visual: two-sample paired t-test for higher activation in FEF stimulation condition (blue) or ACC (pink) compared to control for the group contrast of videos versus black screen (n=4, cluster-corrected at z>2.3, p<0.05 FWE corrected). **b,** Social: two-sample paired t-test for higher activation in FEF stimulation condition (blue) or ACC (pink) compared to control for the group contrast of social videos versus scrambled videos (n=4, cluster-corrected at z>2.3, p<0.05 FWE corrected).

**Supplementary Figure 8.**
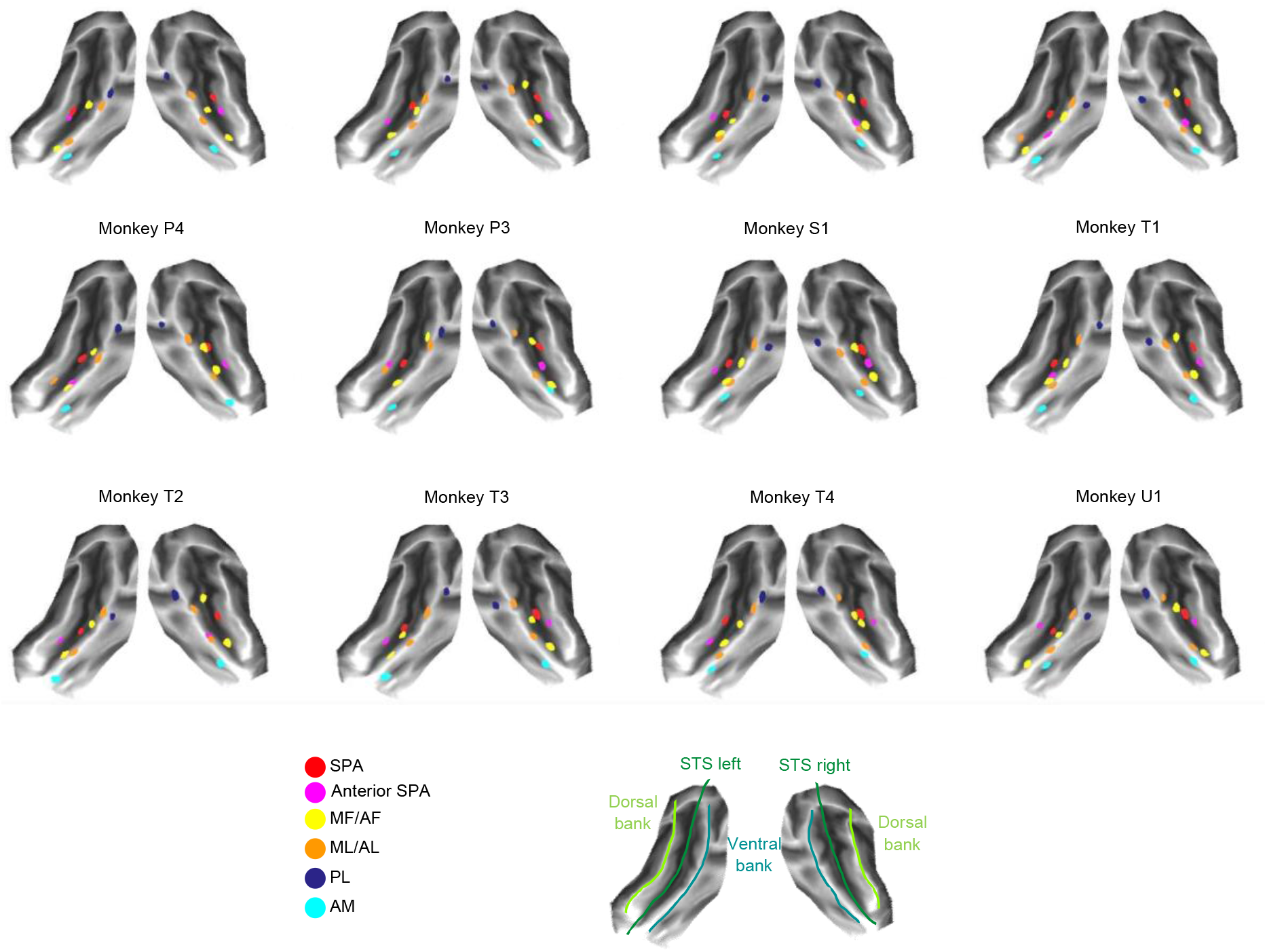
Peak activities for individual macaques. Peak activity for the SPA and for the face patches represented on a flat F99 surface showing the STS with its dorsal and ventral bank.

**Supplementary Table 1.** Peak activation coordinates of social prediction and controls at the group level.

|  | Social prediction |  |  | Visual control |  |  | Object control |  |  |
| --- | --- | --- | --- | --- | --- | --- | --- | --- | --- |
|  | x | y | z | x | y | z | x | y | z |
| midSTS Rostral | 24.6 | -11.6 | -2.51 |  |  |  |  |  |  |
| midSTS caudal | 22.6 | -16.6 | 3.02 | 22.1 | -17.1 | 1.51 | 23.1 | -17.1 | 1.01 |
| FEF | 15.1 | 7.04 | 16.6 |  |  |  |  |  |  |
| V1/V2 | 8.05 | -37.2 | 18.1 |  |  |  |  |  |  |
| V2 lateral | 13.6 | -33.2 | 1.01 |  |  |  |  |  |  |
| V2 medial | 1.51 | -31.2 | 6.54 |  |  |  |  |  |  |
| Superior colliculus | 2.52 | -20.1 | 1.51 |  |  |  |  |  |  |

Coordinates given for the right hemisphere in mm in F99 standard space (Social prediction and visual control: n=14, cluster-corrected at $z>3.1$ and $p<0.05$ FWE corrected, object control: n=7, cluster-corrected at $z>2.3$ and $p<0.05$ FWE corrected).

**Supplementary Table 2.**
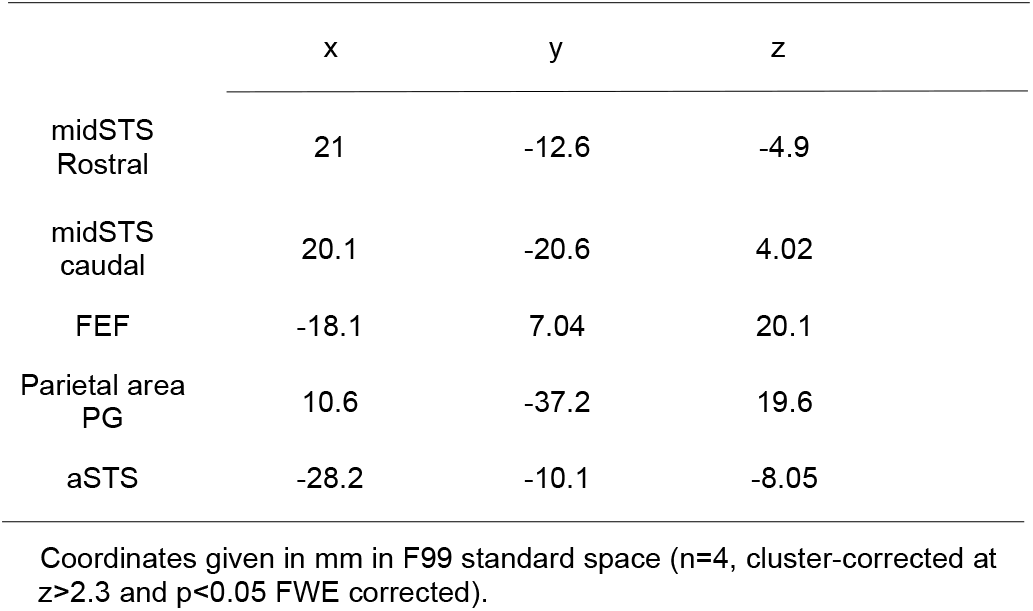
Peak activation coordinates of social prediction at the group level for the replication study.

|  | x | y | z |
| --- | --- | --- | --- |
| midSTS Rostral | 21 | -12.6 | -4.9 |
| midSTS caudal | 20.1 | -20.6 | 4.02 |
| FEF | -18.1 | 7.04 | 20.1 |
| Parietal area PG | 10.6 | -37.2 | 19.6 |
| aSTS | -28.2 | -10.1 | -8.05 |

Coordinates given in mm in F99 standard space (n=4, cluster-corrected at $z>2.3$ and $p<0.05$ FWE corrected).

**Supplementary Table 3.** Peak activation coordinates of the face localizer at the group level.

|  | Right |  |  | Left |  |  |
| --- | --- | --- | --- | --- | --- | --- |
|  | x | y | z | x | y | z |
| AM | 21.1 | -1.5 | -16.3 | -20.5 | -0.7 | -14.7 |
| AF | 22.6 | -2 | -10.1 | -20.5 | -5.2 | -9.8 |
| AL | 24.6 | -0.9 | -9.5 | -22.5 | -1.17 | -10.6 |
| MF | 25.2 | -15.6 | -0.5 | -22.0 | -14.6 | -4.6 |
| ML | 30.7 | -16.1 | 4.5 | -27.6 | -17.2 | 2.5 |
| FEF | 21.1 | 10.6 | 8.6 | -21.1 | 10.6 | 6.54 |

Coordinates given for the right and left hemisphere in mm in F99 standard space (n=14, cluster-corrected at $z>3.1$ and $p<0.05$ FWE corrected).

**Supplementary Table 4.**
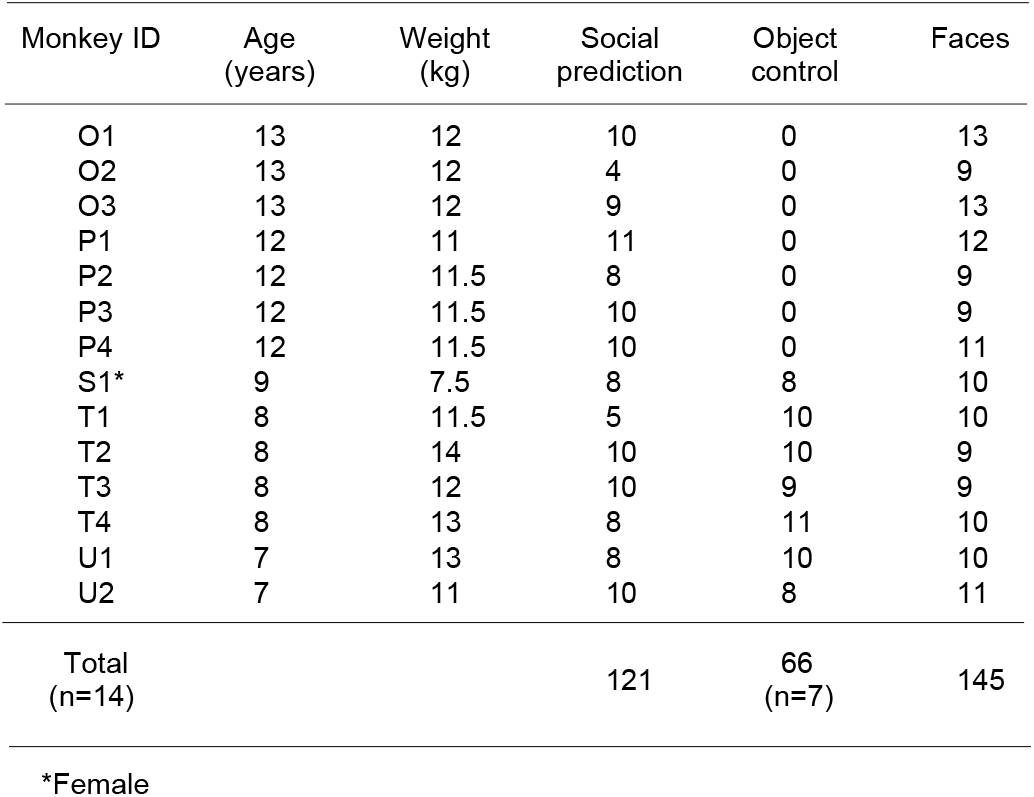
Detail of the monkeys and number of runs per subjects and per conditions selected for analysis.

| Monkey ID | Age<br>(years) | Weight<br>(kg) | Social<br>prediction | Object<br>control | Faces |
| --- | --- | --- | --- | --- | --- |
| O1 | 13 | 12 | 10 | 0 | 13 |
| O2 | 13 | 12 | 4 | 0 | 9 |
| O3 | 13 | 12 | 9 | 0 | 13 |
| P1 | 12 | 11 | 11 | 0 | 12 |
| P2 | 12 | 11.5 | 8 | 0 | 9 |
| P3 | 12 | 11.5 | 10 | 0 | 9 |
| P4 | 12 | 11.5 | 10 | 0 | 11 |
| S1* | 9 | 7.5 | 8 | 8 | 10 |
| T1 | 8 | 11.5 | 5 | 10 | 10 |
| T2 | 8 | 14 | 10 | 10 | 9 |
| T3 | 8 | 12 | 10 | 9 | 9 |
| T4 | 8 | 13 | 8 | 11 | 10 |
| U1 | 7 | 13 | 8 | 10 | 10 |
| U2 | 7 | 11 | 10 | 8 | 11 |
| Total<br>(n=14) |  |  | 121 | 66<br>(n=7) | 145 |

\*Female

